# The Pipeline for Digital Analysis of IHC Images from NSCLC Xenograft Tissues

**DOI:** 10.1101/826545

**Authors:** Rachana Kandru, Bina Desai

## Abstract

Multiple small molecule inhibitors and immunotherapy advances have shown success by prolonging patient’s survival in NSCLC but patient with metastasis or advanced stage of cancer often experiences relapse. Cancer associated fibroblast (CAFs), a major cellular component of tumor microenvironment plays key role in shaping solid tumors. *In-vitro* studies have reported that CAFs can blunt the effect of targeted therapies in tumor cells. The current study focuses on evaluating the impact of stromal protection by analyzing immunohistochemistry (IHC) images from xenograft experiment. The study investigates if the HGF-driven CAF mediated stromal protection in cancer cell contributes to the development of resistance to Alectinib. We utilized QuPath, a digital software to automatize the readout of proliferation rate of tumor cells and evaluate the impact of stromal protection.

## 1. Background

Lung cancer is the second most common cancer in both men and women^1^. Recent advances with Tyrosine kinase inhibitors (TKIs) and immunotherapies have shown success in prolonging patient’s survival but the resistance develops inevitably in advance and metastatic non-small cell lung cancer (NSCLC). Tumor, as we have discovered, is not only composed of tumor cells, but rather has complex microenvironment where tumor and the surrounding microenvironment are closely related and constantly interact with each other. Tumor ecosystem includes blood vessels, immune cells, signaling molecules, extracellular matrix, and fibroblasts. Each of these components make their own respective contribution to either pro-tumorigenic or anti-tumorigenic activity. In case of Anaplastic Lymphoma Kinase positive (ALK+) NSCLC, most of the tumor cells are ruled out when patients are treated with ALK inhibitors leaving behind small population of tumor cells which remains unaffected and contributes towards development of resistance. The ability of tumor cells to survive under therapy pressure is well documented by their intrinsic capabilities^2,3^ or microenvironmental factors such as pro-survival signals which promote the resistance phenotype in cancer cells^4^.

Cancer Associated Fibroblasts (CAFs) play a key role in augmenting the tumorigenic potential of cancer cells. Multiple in-vitro studies reported that CAFs promote tumor progression^5,6^ CAFs contributes in shaping the tumor structure by releasing growth factors and paracrine signaling factors which stimulates tumor growth. These pro-signaling factors are reported the blunt the effect of therapeutic response in cancer cells^7,8^. Our study focuses on evaluating the impact of CAF-mediated stromal protection on evolution of resistance. The study will elaborate if the CAF proximity can impact the evolution of resistance in xenograft mouse model.

### 1. CAFs blunts the effect of Alectinib (*in-vitro*)

We used H3122 cells and cultured them with conditioned media (CM) from human CAFs. The cells were treated with either DMSO vehicle or Alectinib. We observed that the cells treated with Alectinib and cultured in CM showed significant increase in cell proliferation rate as compared to cells treated with Alectinib & cultured without conditioned media

### 2. Mouse Fibroblasts do not protect tumor cells

To recapitulate the results *in-vivo*, we isolated mouse embryonic fibroblasts (EF). H3122 cells either co-cultured with EF or with CM from EF and cells treated with 0.25µM or 0.5µM of Alectinib or DMSO. The results indicated that mice fibroblasts were not able to protect H3122 when treated with Alectinib (Fig. 1). This rose a new question: what is the difference between human CAFs and mouse CAFs?

**Figure 1.**
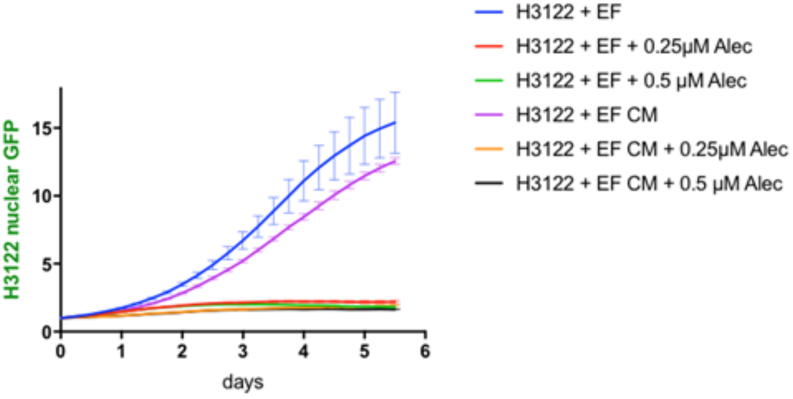
Mouse embryonic fibroblasts do not protect H3122 cells when treated with Alectinib

### 3. HGF an important contributor but not sufficient for CAF mediated protection

In human embryonic organs, Hepatocyte growth factor (HGF) is mainly produced in stroma cells such as fibroblasts, and is important for tissue repair and protection in vivo. By serving as a paracrine regulator for embryogenesis and organ regeneration, HGF plays a key role in tumor protection. Multiple studies have reported the importance of HGF-mediated CAF stromal protection^7,9,10^. In our ALK+ NSCLC experimental model, we found HGF is required for CAF-mediated stromal protection but is not sufficient to promote the tumor growth (Fig. 2).

**Figure 2.**
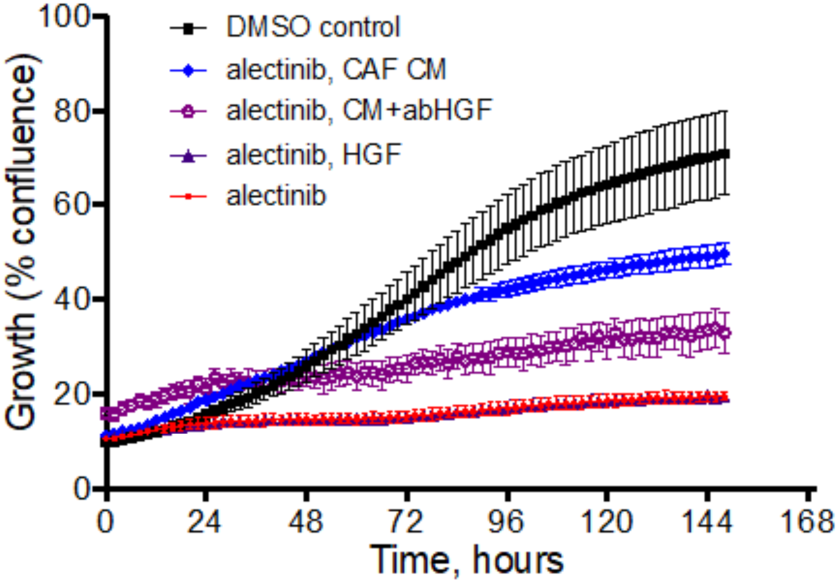
HGF contributes to CAF-mediated stromal protection

### 4. Mice Model

Since murine HGF cannot activate human cMET, we acquired humanized HGF NSG (human allele for HGF instead of a mouse allele, hHGF) mice. In our mouse experiment, we utilized 5 of each NSG and hHGF NSG mice, both the groups were injected with H3122 cells via subcutaneously and were either treated with Alectinib (25 mg/kg, daily, oral gavages) or left untreated (control group). Animals were sacrificed if tumors grew more than 2mm^2^. Tumor tissues were extracted and stained for BrdU+ (proliferation marker). By looking at the BrdU+ from NSG and hHGF mice. This experimental will address if the effect of stromal protection in NSG versus hHGF mouse in response to Alectinib. BrdU+ cells will serve as experimental read out, implementing the stromal protection could promote proliferation of tumor cells. In parallel, 5 of NSG mice were utilized and H3122 cells were introduced via sub-cutaneous injection in each mouse. These mice were sacrificed at either initial time point, after short term treatment (4 cycles of Alectinib, 25mg/kg) and at the late stage of continuous treatment (relapse). Tumor tissues were extracted and stained for BrdU+ marker.

### 5. Methodology: Image Processing

QuPath was used to analyze the IHC slides from both mice models. QuPath enabled us to create neural network that which can detect BrdU+ cells, stroma and tumor cells by training multiple sections on each slide. In order to reach closer to accuracy, we trained the software with approximately 200 manual segmentations (tumor/stroma/BrdU+ cells, excluding necrotic tissue) to create neural network. The trained neural network was used to analyze multiple sections from each slide, followed by manual quality control check to avoid any error in detecting different types of cells.

### 6. Results

Immunohistochemistry stain showed both NSG and hHGF NSG mice when treated Alectinib showed significant decrease in %BrdU cells (Fig.4) but there was no different in within both mouse models. Suggesting, HGF driven CAF-mediated stromal effect might be too subtle to see the difference by analyzing tumor volume.

**Figure 3.**
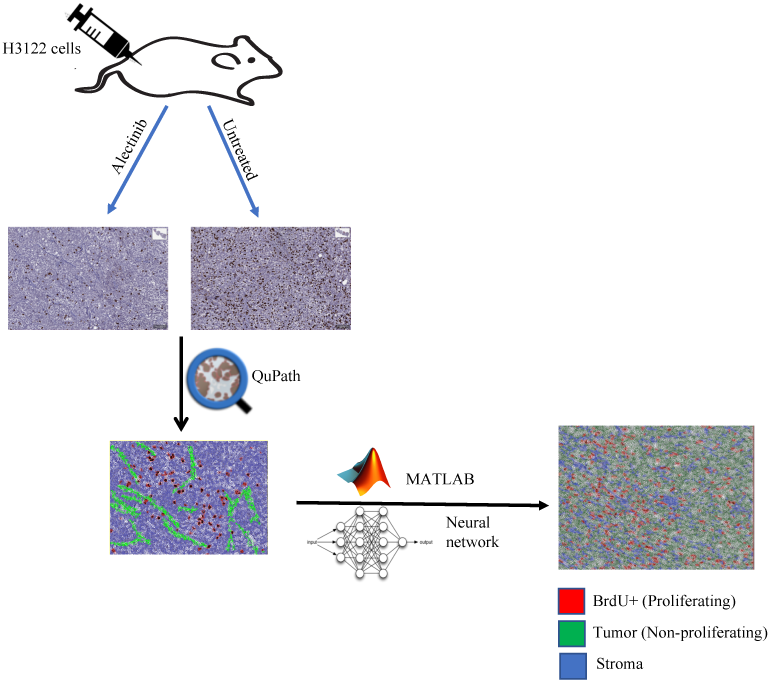
Workflow for digital analysis from xenograft tissues.

**Figure 4.**
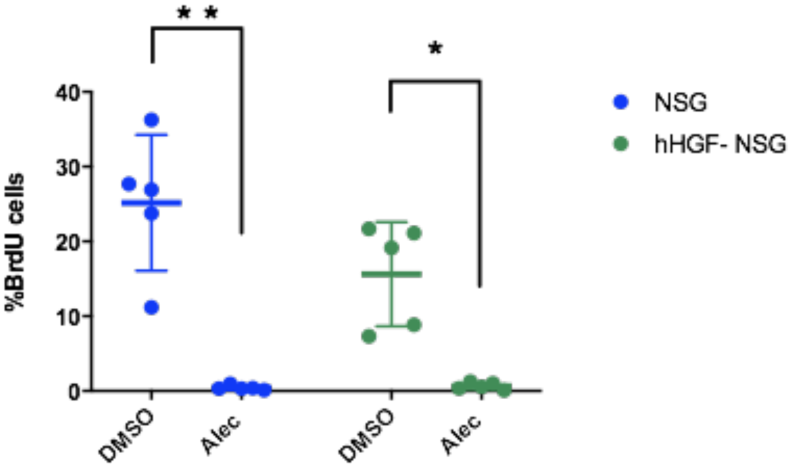
Comparing BrdU+ positive cells in NSG versus hHGF-NSG mice (**p<0.0001, *p<0.005)

From our second animal study we observed significant decrease in %BrdU+ cells with the short-term treatment but the %BrdU counts increases as the treatment is extended (Fig.5), confirming the relapse phenomenon where tumor starts proliferating while developing resistance to Alectinib.

**Figure 5.**
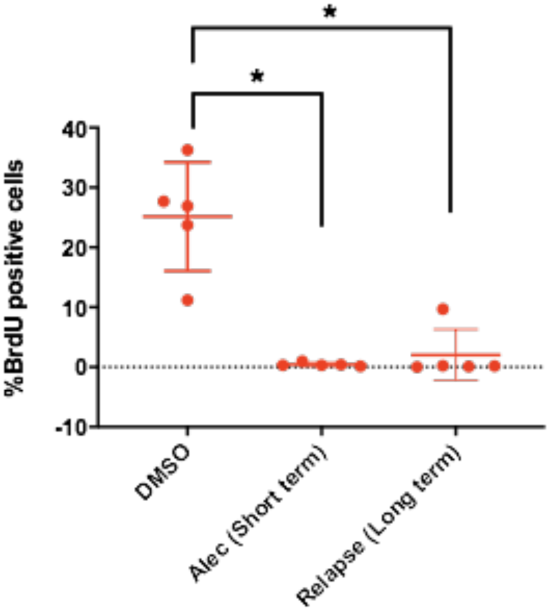
Comparing BrdU+ positive cells in control and Alectinib (short term and long term/relapse) NSG mice, (one-way ANOVA p< 0.0001)

### 7. Conclusion

Our study shows the digital analysis of IHC slides can quantitatively extract information from 2D-slides. From both the mouse model experiment, our digital analyses showed decrease in %BrdU+ cells when mice were treated with Alectinib, which is expected as Alectinib treatment will kill most of tumor cells. While we didn’t observe any difference in response to Alectinib treatment with NSG and hHGF NSG mice, we believe further investigation is required to dissect the role of HGF-driven stromal protection.

### 8. Future Steps

We plan to utilize our digital platform to analyze multiple slides from NSG and hHGF-NSG mice to study if proximity of CAF to tumor cells can trigger the stromal protection as compared to tumor cells residing distant to CAF. Additionally, we observed that short term Alectinib treated mice showed decrease in %BrdU+ cells, but we don’t know if this is because the cells are no longer proliferating or a greater number of cells are dying? To address this question, we plan to stain slides with Cleaved caspase 3 to analyze the % apoptotic cells in each group. Further, all of the above data will be utilized to generate math model to predict the impact of stromal protection on evolution of resistance.

## Acknowledgments

B.D performed *in-vitro* experiments and R.K analyzed the data using QuPath software. R.K. and B.D would like to thank Olena Balynska for technical assistance, Dr. Heiko Enderling, and Mark Laurie for their valuable help and Dr. Andriy Marusyk for performing *in-vivo* experiments & critical reading of this article.

